# The vasodilator-stimulated phosphoprotein (VASP) supports the conduction of vasodilator signals and NO-induced arteriolar dilations in murine arterioles *in vivo*

**DOI:** 10.1101/2023.03.17.533095

**Authors:** Philip Poley, Peter M. Benz, Cor de Wit

## Abstract

VASP is a member of the Enabled/VASP protein family that is involved in cortical actin dynamics and may also contribute to the formation of gap junctions. In vessels, gap junctional coupling allows the transfer of signals along the vessel wall and coordinates vascular behavior. Moreover, VASP is reportedly a mediator of NO-induced inhibition of platelet aggregation. Therefore, we hypothesized that VASP exerts also important physiologic functions in arterioles. We examined the spread of vasodilations enabled by gap junctional coupling in endothelial cells as well as NO-induced arteriolar dilations in VASP-deficient mice by intravital microscopy of the microcirculation in a skeletal muscle in anesthetized mice. Conducted dilations were initiated by brief, locally confined stimulation of the arterioles with acetylcholine. The maximal diameters of the arterioles under study ranged from 30 to 40 μm. Brief stimulation with acetylcholine induced a short dilation at the local site that was also observed at remote, upstream sites without an attenuation of the amplitude up to a distance of 1.2 mm in control animals (wild-type). In contrast, remote dilations were reduced in VASP-deficient mice despite a similar local dilation indicating an impairment of conducted dilations. Superfusion of NO-donors induced a concentration-dependent dilation in wild-type mice. However, these dilations were slightly reduced in VASP-deficient animals. In contrast, dilations induced by the endothelial stimulator acetylcholine were fully preserved in VASP-deficient mice. In summary, this study suggests that VASP exerts critical functions in arteriolar diameter control. It is crucial for the conduction of dilator signals along the endothelial cell layer. The impairment possibly reflects a perturbed formation of gap junctions in the endothelial cell membrane. VASP also participates in the full dilatory potential of NO-donors although the effect of its deficiency is only subtle. In contrast, VASP is not required for dilations initiated by endothelial stimulation which are mediated in the murine microcirculation by an EDH-mechanism.

## Introduction

Vasodilator-stimulated phosphoprotein (VASP) is a member of the Enabled/VASP protein family. These proteins are involved in cortical actin dynamics. Cortical actin assembly is a driving force for intercellular adhesion and the assembly of actin is regulated by alphaII-spectrin-VASP complexes in endothelial cells (Benz et al., 2008). The actin cytoskeleton is also crucial in the transport of gap junctional proteins to the membrane (Smyth et al., 2012; Vanslyke et al., 2009) and the formation of functional gap junctions within the membrane (Thevenin et al., 2013). Therefore, VASP may also contribute crucially in gap junction formation in the vessel wall.

Cellular coupling and signal transmission via gap junctions is of functional importance in vascular tissue. Endothelial and smooth muscle cells in the vessel wall are coupled heterocellularly via gap junctions thereby providing a signaling pathway in the transversal direction, i.e. from endothelial to smooth muscle cells and vice versa (myoendothelial coupling). This coupling enables the transfer of an endothelial cell hyperpolarization into the adjacent smooth muscle cells that relax upon hyperpolarization. Thereby, myoendothelial coupling underlies the relaxation upon endothelium-dependent hyperpolarization (EDH-dilation) although other mechanisms may also contribute (Schmidt and de Wit, 2020; Shu et al., 2019; Straub et al., 2014). In addition, endothelial cells and vascular smooth muscle cells are coupled homocellularly thereby providing a longitudinal signaling pathway along the vessel. This signaling pathway serves to coordinate the cellular behavior in arterioles in the microcirculation along the vessel length by integrating cells into a tightly coupled syncytium creating a large cell cluster (Schmidt et al., 2008). It was suggested that this pathway contributes to the matching of oxygen supply and tissue needs (Bagher and Segal, 2011). In fact, under conditions of enhanced tissue oxygen needs (skeletal muscle contraction) mice with impaired gap junctional communication exhibit reduced dilations indicating the physiologic importance of this pathway in active hyperemic responses induced by metabolic demand (Milkau et al., 2010).

The longitudinal conduction pathway is mainly provided by endothelial cells because they are tightly coupled through gap junctions composed of Cx40 and Cx37. Of these connexins, Cx40 is of outmost importance and deletion of Cx40 impaired longitudinal signaling (de Wit et al., 2000). In addition, the lack of Cx40 reduced the expression of Cx37 in endothelial cell membranes suggesting that Cx40 is required to transfer Cx37 into the cell membrane (Jobs et al., 2012). Functionally, the tight coupling is reflected by the spread of a locally induced vasomotor response along the vessel which is called a conducted response. Mechanistically, it was demonstrated that an electrotonic spread of a locally initiated hyperpolarisation underlies the conduction of the dilation. Interestingly, the hyperpolarisation is amplified in the endothelium while the signal spreads along the vessel and therefore conducted dilations cover considerable distances in the microcirculation (Jackson, 2017).

Reportedly, platelet adhesion is enhanced in VASP-deficient mice and the inhibitory effect of NO on platelet adhesion was abrogated in these animals indicating that VASP mediates the inhibitory effect of exogenous cGMP-elevating compounds as well as being involved in endogenously derived NO inhibitory effects in platelets (Massberg et al., 2004). However, VASP was not required for the inhibition of platelet aggregation by the P2Y12 receptor antagonist in rats (Ito et al., 2018). Vascular smooth muscle cell relaxation is likewise induced by substances that elevate the second messengers cAMP or cGMP in arterioles in the microcirculation (de Wit et al., 1994) and therefore VASP may also be invoked in smooth muscle relaxation. Accordingly, VASP phosphorylation was enhanced in smooth muscle cells in rat aorta by cAMP- and cGMP-elevating compounds (Schafer et al., 2003). However, relaxation of the aorta directly induced by cGMP or cAMP remained unaffected by VASP deletion although the inhibition by cAMP and cGMP on platelets was abrogated in these VASP-deficient mice (Aszodi et al., 1999). On the other hand, VASP phosphorylation was required to induce effects of NO on smooth muscle cell adhesion and spreading (Defawe et al., 2010).

## Material and methods

### Animals

Animal care and experiments were in accordance with the German Animal Welfare Act and approved by local authorities (Ministerium für Landwirtschaft, Umwelt und ländliche Räume des Landes Schleswig-Holstein). VASP-deficient mice (Benz et al., 2008; Hauser et al., 1999; Laban et al., 2018) and their respective WT littermates were bred at the Goethe University Hospital animal facility. We examined male VASP-deficient mice and their wildtype littermates as controls.

### Intravital microscopy of the microcirculation

Mice were anesthetized with intraperitoneal injection of fentanyl (0.05 mg/kg), midazolam (5 mg/kg), and medetomidin (0.5 mg/kg) in a volume of 13 mL/kg. A jugular vein catheter allowed subsequent continuous infusion of the anesthetic drugs (at 5 mL/kg/h and 0.02, 2.2, 0.22 mg/kg/h of fentanyl, midazolam, and medetomidin). The infusion rate was adjusted according to the needs as checked by whisker and paw reflexes. A small tube was inserted into the trachea to secure the airway and to ventilate the animal during the experiment using ambient air with a tidal volume of 200 μl at a rate of 150 /min (MicroVent Mouse Ventilator, Hugo Sachs Elektronik, Hugstetten, Germany). Throughout the preparation and during the experiment, mice were warmed by an electronically heated mat adjusted to 37°C. The cremaster muscle was prepared as described (Jobs et al., 2012) superfused at 8 mL/min with a tempered (34° C) buffer containing (in mmol/L): 118.4 NaCl, 20 NaHCO_3_, 3.8 KCl, 2.5 CaCl_2_, 1.2 KH_2_PO_4_, 1.2 MgSO_4_. The buffered (pH 7.4) saline solution was gassed with 5% CO_2_ and 95% N_2_ resulting in a pO_2_ of about 30 mmHg on the cremaster muscle due to contamination with ambient air. Arterioles were observed through a 20- or 40-fold objective using a video camera-equipped microscope (Axioscope FS, Zeiss, Germany). Images were recorded on videotape to allow later measurement of the inner arteriolar diameter with an application written in LabVIEW8.0 (National Instruments, Austin, Texas, USA). This application projected a rectangle over digitized images which boundaries marked the inner arteriolar diameter. For diameter measurements, the user adjusted the boundaries of this rectangle by simple key strokes during replay of the microscopic images at reduced speed. Values were sampled at a frequency of 2 Hz. A series of different arterioles was measured by drawing a rectangle that was projected over the still images of single arterioles and aligned to the inner arteriolar diameter by the user using the computer mouse.

### Experimental procedure

After 30 minutes of equilibration the preparation was inspected and arterioles were chosen for examination. Criteria for the selection of arterioles were clear visibility, branching order, length, and accessibility for stimulation using glass pipettes.

For studying conducted vasodilations, a glass micropipette filled with acetylcholine (ACh, 1 mmol/L) was positioned in close proximity to a second- or third-order arteriole as described (Jobs et al., 2012). In brief, a short pressure pulse (150 kPa, 100 to 500 ms) applied by a pneumatic ejector (PDES 02D, npi, Germany) onto the micropipette led to a locally confined stimulation of the arteriole while the microscopic images and the arteriolar response was recorded. Recording was started 30 s before and lasted until 90 s after the stimulation. If a response at the site of stimulation (local) was observed, the microscope was moved to upstream sites at a distance of 0.6 mm or 1.2 mm and the stimulation was repeated without interfering with the stimulation pipette. The pressure pulse was applied again and the response at an upstream site was recorded. After examining local and two remote sites in duplicate, another arteriole was chosen and the stimulation pipette was positioned in the proximity of this arteriole. Up to three arterioles were studied in a single experimental that lasted up to 4 hours.

In a second protocol, we examined concentration-response-curves for different vasoactive agonists that were added to the superfusion fluid thereby accessing the complete preparation and each arteriole within. In each mouse 8-12 arterioles were chosen that were clearly visible. Arteriolar diameters were measured before and during addition of ACh, adenosine or the NO-donors sodium-nitroprusside (SNP) or DEA-NONOate [2-(N,N-Diethylamino)-diazenolate-2-oxide] to the superfusion buffer. Before studying the next concentration or the next vasodilator, arterioles were allowed to return to their resting diameter for 4 min. By this means, non-cumulative concentration response curves were generated.

In all experiments, the maximal luminal diameter of all arterioles under study was measured in the presence of a maximally dilating stimulus (combined superfusion of sodium nitroprusside, acetylcholine, and adenosine; each 30 μmol/L). Experiments typically lasted 2 to 4 hours. At the end of the experiment, animals were sacrificed by an overdose of pentobarbital (2.5 g/kg) applied intravenously.

### Data analysis

Vascular tone is given as percentage of maximal diameter. Internal diameters were measured and changes normalized to the dilator capacity according to the formula:

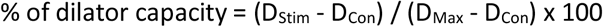

with D_Stim_ being diameter during stimulation, D_Con_ control diameter before stimulation, and D_Max_ the maximal diameter measured for the arteriole during the experiment which included the combined superfusion of different dilators at supramaximal concentrations. Statistical analysis was performed using STATA (Stata Corporation Texas, USA). Data are presented as a mean ± standard error of mean (mean ± S.E.M.). Data between groups (VASP-deficient vs. wildtype mice) were analysed by unpaired t-test. Paired samples (arterioles at different sites during the study of conducted dilations) were analysed by paired t-test. Differences were considered significant at an error probability of *P* < 0.05.

## Results

### 1. VASP and gap junctional coupling in the vessel wall

Arterioles under study exhibited maximal diameters between 30 and 40 μm. The arteriolar resting diameters were similar in both genotypes and ranged from 10 and 15 μm (wildtype: 11.3±1.4 μm, VASP-deficient: 13.0±1.2 μm, not significantly different from each other). Brief, locally confined stimulation by short pressure-induced ejection of acetylcholine from a micropipette positioned in the close vicinity of the arteriole led to a rapid and large dilation at the stimulation site (Fig. 1A) that lasted for about 30s. This dilation conducted without measurable delay along the arteriole and was observed also at remote upstream sites in a distance of 0.6 and 1.2 mm (Fig. 1B, Fig. 1C). The amplitude of the dilation at remote sites was not reduced compared to the local stimulation site in wildtype animals (Fig. 1D). The dilation upon acetylcholine stimulation was of a similar amplitude in VASP-deficient mice at the local stimulation site (Fig. 1A) and also conducted to remote sites without delay (Fig. 1B, Fig. 1C). However, in contrast to wildtype animals, the amplitude of the dilation was significantly reduced in VASP-deficient mice at remote sites (Fig. 1D). This attenuation of the amplitude of conducted dilations resembles the observation in mice deficient for Cx40 in endothelial cells (de Wit et al., 2000; Jobs et al., 2012). We also examined connexin expression in endothelial cells in these arterioles by immunostaining. However, we did not find an obvious reduction or misassembly of Cx40 or Cx37 in VASP-deficient arterioles (data not shown).

**Figure 1:**
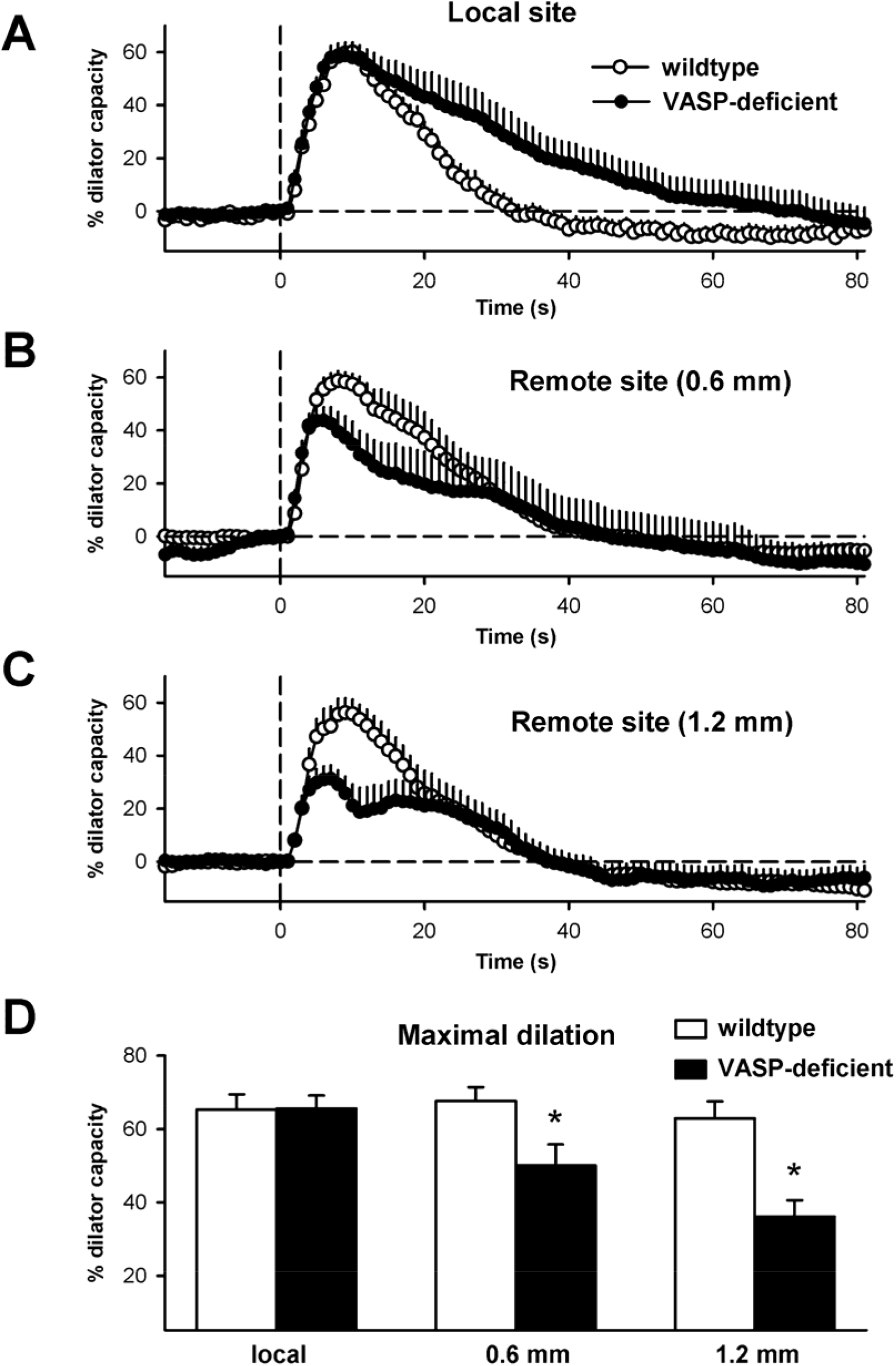
Arterioles in the skeletal muscle microcirculation of mice were briefly stimulated in a locally confined manner with acetylcholine to examine conducted dilations. Arteriolar diameter changes are plotted as % of dilator capacity over time at the stimulation site (A) and upstream, remote sites in a distance of 0.6 mm (B) and 1.2 mm (C). The stimulation with acetylcholine (at time point 0, dashed line) induced a rapid dilation at the stimulation site (A, local site) in wildtype (open symbols) and VASP-deficient mice (black symbols). The dilation conducted without measurable delay along the vessel and was also observed at upstream remote sites (B, 0.6 mm; C, 1.2 mm) in both genotypes. In (D) the maximal amplitude during the observation period is depicted for the different sites studued. Whereas in wildtype mice (open bars) the dilatory amplitude did not decrease up to a distance of 1.2 mm, the amplitude of the dilation was significantly attenuated at remote sites (0.6 and 1.2 mm) in VASP-deficient mice (black bars). Experiments were performed in 5 wildtype and 4 VASP-deficient animals, in each animal up to three arterioles were studied, data represent n=14 or 12 arterioles; * indicates *P*<0.05 vs. local dilation (paired t-test, Bonferroni corrected).

### 2. Role of VASP in arteriolar dilation

We examined the role of VASP in arteriolar dilations in the murine microcirculation by studying concentration-dependent dilator responses in VASP deficient mice. For each genotype (wildtype, VASP-deficient), 65 arterioles were studied in 5 mice. The resting and maximal diameters of the arterioles under study were not different between genotypes (resting diameter: 8.9±0.5 vs. 9.1±0.5 μm; maximal diameter: 28.2±1.0 vs. 29.7±0.8 μm, wildtype and VASP-deficient, respectively). In these arterioles, the application of NO-donors (sodium nitroprusside [SNP] or DEA-NONOate) induced concentration-dependent dilations. However, these dilations were reduced in VASP-deficient arterioles at intermediate concentrations of SNP (by about 25%, Fig 2A) and at the highest concentration of DEA-NONOate (by about 13%, Fig 2C). Thus, a substantial dilation remained intact in the absence of VASP indicating that VASP is not a critical mediator of smooth muscle relaxation in murine arterioles. Nevertheless, the dilator potential of NO was attenuated in VASP-deficient arterioles suggesting that VASP contributes in the dilatory effect of NO. Since the dilations were only partially reduced and this was only observed at some concentrations, these observations are in line with a rightward shift of the concentration response curve (NO desensitization). Similarly, a different dilator (adenosine) which acts mainly via elevation of cAMP and opening of potassium channels (de Wit, 2010) was likewise slightly reduced at a medium concentration (Fig 2D). In marked contrast, dilations induced by the endothelium-dependent agonist acetylcholine were fully preserved (Fig 2B).

**Figure 2:**
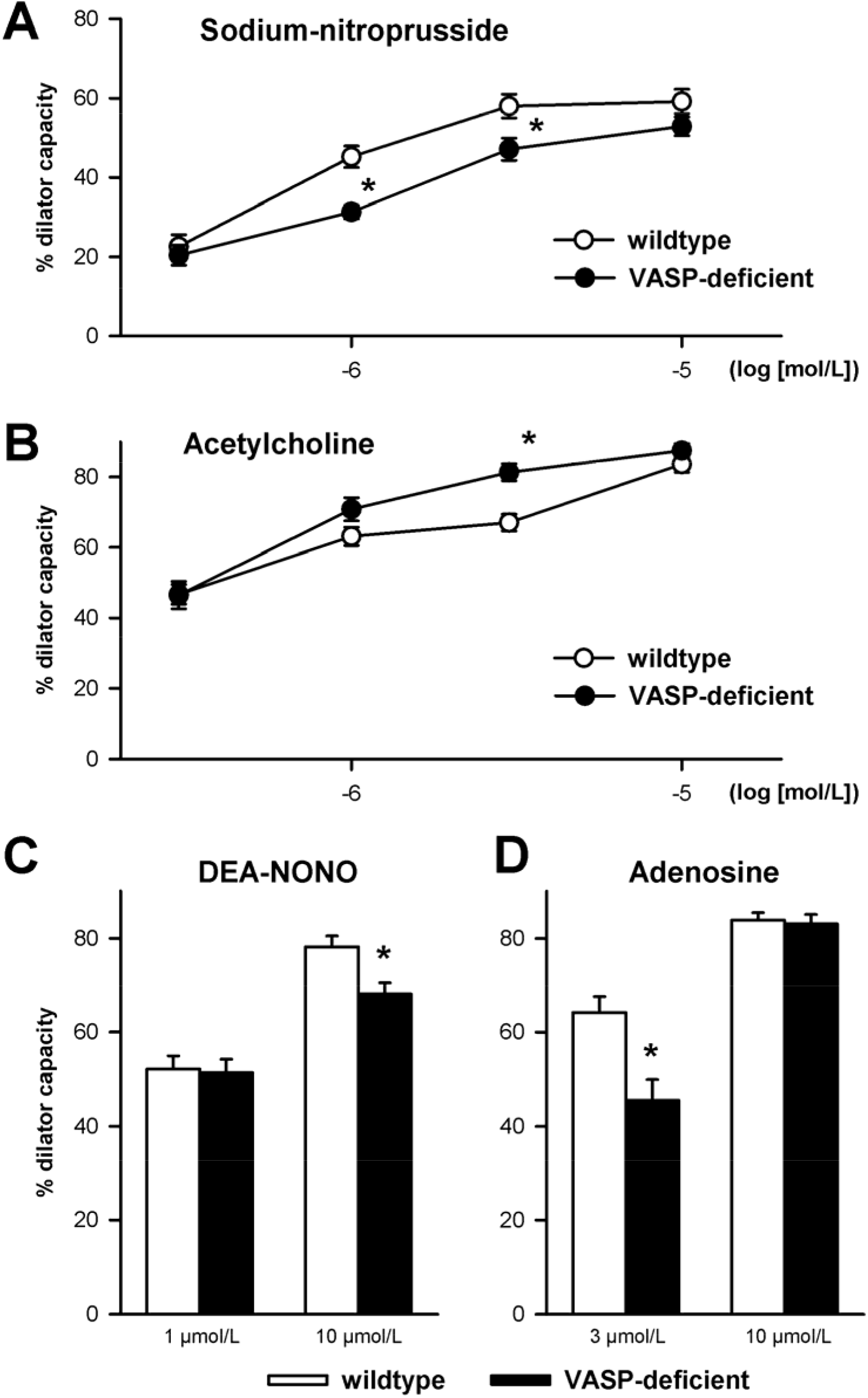
Arterioles in the skeletal muscle microcirculation of wildtype (white symbols) and VASP-deficient mice (black symbols) were monitored before and during application of different vasoactive substances, the NO-donors sodium-nitroprusside (A) or DEA-NONOate (C), the endothelial agonist acetylcholine (B), and the cAMP-elevating agonist adenosine (D). Dilation is depicted as % of dilator capacity. The NO-donors (A, C) induced a concentration-dependent dilation in wildtype mice. NO-induced dilations were also found in VASP-deficient mice, but the responses were reduced indicating that VASP is not critical for NO-induced cGMP-mediated dilations but it supports the dilation. The data indicate a rightward shift of the concentration response curve. A similar, small attenuation was also present for the cAMP-elevating agonist adenosine (D). In contrast, endothelial stimulation using acetylcholine and the subsequent EDH-type dilation was fully unhindered (or even slightly enhanced) in VASP-deficient mice. Experiments were performed in 5 mice of each genotype, in each animal 13 arterioles were studied, data represent n=65 arterioles for each genotype; * indicates *P*<0.05 vs. wildtype (unpaired t-test).

## Discussion

This study suggests that VASP exerts crucial functions in arteriolar diameter control. However, the effects observed in the murine microcirculation were only subtle. The spread of membrane potential changes along the vascular wall through gap junctions support the coordination of cellular behavior along the vessel length that yields into a dilation encompassing considerable distances of arterioles in the microcirculation. Such behavior can be experimentally tested by locally confined stimulation with agonists that induce a membrane potential change in endothelial cells such as acetylcholine (de Wit, 2010). Our data suggest that VASP exerts a critical role in supporting the spread of hyperpolarisations along the endothelial cell layer and consequently in the conduction of dilations along the vessel wall as indicated by the reduced amplitude of the dilation at remote sites. Indeed, the propagation of electrical signals is also delayed in cardiac tissue in Ena/VASP-deficient animals most likely resulting from impaired gap junction assembly (Benz et al., 2013). In cardiac ventricles, Cx43 is the major isoform found and is the most important connexin supporting gap junctional coupling although Cx43 levels have to decline considerably before a decrease of the conduction velocity becomes evident (Danik et al., 2004; Lambiase and Tinker, 2015). In vessels, Cx40 is crucial to support the propagation of electrical signals between endothelial cells and thereby enable conducted dilations in arterioles (de Wit et al., 2000). Despite the different isoforms being involved, we suggest that VASP is also important for the proper recruitment of connexins into the cell membrane and the composition of functional gap junctions in the endothelium of the arteriolar wall. However, we were unable to verify a reduced expression of connexin40 in the endothelial cell membrane that is most likely due to the vague quantification of protein expression possible by immunostaining.

In addition, we suggest that VASP contributes to the full dilatory potency of exogenous NO applied onto the microcirculation. Importantly, VASP is not a requirement because a considerable portion of the NO-induced dilation remained in VASP-deficient mice. The impairment of the dilation was observed at intermediate NO-donor concentrations implying a shift of the concentration-response curve towards higher concentrations of these cGMP-elevating compounds. In marked contrast, dilations to the endothelial against acetylcholine remained fully intact. It has to be noted that acetylcholine dilates arterioles in the murine microcirculation through an EDH-mechanism and is not mediated by endothelial NO release (Boettcher and de Wit, 2011; Schmidt and de Wit, 2020; Si et al., 2006)230003, 180485, 320053). Therefore, we conclude that VASP is not a crucial actor in arteriolar dilations in general, but only supports dilations in response to vasodilators that act by elevation of the second messenger cGMP in the vascular smooth muscle cell. In contrast to NO-donors, EDH-dilations which act by inducing smooth muscle hyperpolarisation are completely independent of VASP.

## Acknowledgements

This work was supported by the German Center for Cardiovascular Research (DZHK B14-028 SE to CdW and PMB). The authors are indebted to Lea Ulrich for scientific input and help with the VASP-deficient mice.

